# *Limnospira* (Cyanobacteria) chemical fingerprint reveals local molecular adaptation

**DOI:** 10.1101/2024.04.19.590274

**Authors:** Théotime Roussel, Cédric Hubas, Sébastien Halary, Mathias Chynel, Charlotte Duval, Jean-Paul Cadoret, Tarik Meziane, Léa Vernès, Claude Yéprémian, Cécile Bernard, Benjamin Marie

## Abstract

*Limnospira* can colonize a wide variety of environments (*e.g.,* freshwater, brackish, alkaline or alkaline-saline water) and develop dominant and even permanent blooms that limit over-shadowed adjacent phototrophs diversity, especially in alkaline and saline environments. Previous phylogenomic analysis of *Limnospira* allowed us to distinguish two major phylogenetic clades (I and II) but failed to clearly segregate strains according to their respective habitats in terms of salinity or biogeography. In the present work, we attempt to determine whether *Limnospira* displays metabolic signatures specific to its different habitats, particularly brackish or alkaline-saline ecosystems, and question the impact of accessory gene repertoires on respective chemical adaptations.

The study of the metabolomic diversity of 93 strains of *Limnospira* from the Paris Museum Collection, grown under standardized lab culture conditions, showed clearly distinct chemical fingerprints that were correlated with the respective biogeographic origins of the strains. The molecules that most distinguish the different *Limnospira* geographic groups are sugars, lipids, peptides, photosynthetic pigments, and antioxidant molecules. Interestingly, these molecule enrichments might represent adaptation traits to the local conditions encountered in their respective sampling environments concerning salinity, light and oxidative stress. We hypothesize that within extreme environments, such as those colonized by *Limnospira*, a large set of flexible genes can provide remarkable adaptation to specific local environmental conditions (*e.g.,* salinity, light, and oxidative pressure). Thus, the occurrence within *Limnospira* population genomes of a specific set of flexible genes potentially involved in the production of certain metabolites may provide valuable adaptative traits that may support the bloom persistence beyond environmental condition variations.

**Importance:** *Limnospira* are ubiquitous cyanobacteria able to colonize and dominate a wide range of alkaline-saline environments around the world according to remarkable adaptative strategies. Phylogenomic analysis of *Limnospira platensis* allowed to distinguish two major phylogenetic clades (I and II) but failed to clearly segregate strains following their habitats in terms of salinity or biogeography. One can presume that the genes found within this variable portion of the genome of these clades could be involved in *L. platensis* adaptation to local environmental conditions. In the present paper, we attempt to determine whether *Limnospira platensis* displays metabolic signatures specific to its different habitats, particularly brackish or alkalinesaline ecosystems, and question the impact of accessory gene repertoire on respective chemical adaptation.

## Introduction

Certain phytoplanktonic organisms can durably colonize their environment. This capability appears to be balanced by their specific ability to perform efficient primary production, to avoid being overshadowed by competitors, and to deal with physico-chemical environmental constraints. This adaptive power is strongly dependent on the energetic cost they must pay for it, in terms of genomic or physiological expenses^1^.

Cyanobacteria are a group of photoautotrophic microorganisms that have colonized a wide range of environments around the world^2–3^. The large diversity of cyanobacteria species is accompanied by a great diversity of primary and/or secondary/accessory metabolites, many of which presenting noticeable biological activities of central interest in key cellular or ecological functions^4–5^. The genus *Limnospira* (anc. *Arthrospira*), better known under its commercial/vernacular name ‘spirulina’, is one of the most extensively cyanobacteria taxa studied so far (*e.g.* more than 300 publications per year since 2020 on PubMed), especially due to its industrial potential (*e.g.* agri-food, pharmaceuticals, cosmetics)^6–7^. Indeed, cultivated *Limnospira* present a remarkably high growth rate under appropriate culture conditions (more than one division per day)^8–9^, and are very rich in proteins (up to 70% of the dry weight)^10^, unsaturated fatty acids^11^, vitamins (at least 9)^12–13^, and various molecules with antioxidant properties (*e.g.* polyphenols, carotenoids, ergothioneine, and biopterins)^12,14^. To this aim, the physiological aptitudes of certain *Limnospira* strains have been examined (*i.e.,* biomass or molecule production affected by modifying culture parameters)^8,15–18^, whereas only few studies have focused on the global ecological trait of this taxon^19^.

In a recent study based on a polyphasic approach combined with comparative genomics, we showed that the *Limnospira* genus is monospecific, represented by one single species, *Limnospira platensis*, which is cosmopolitan^20^. Yet, *Limnospira platensis* seems to develop a singular ecological strategy, as this species is able to colonize a wide variety of environments (*e.g.*, freshwater, brackish, alkaline or alkaline-saline water)^19–20^ in which it dominates the phytoplanktonic community by developing permanent blooms that limit overshadowed adjacent phototroph diversity^19,21^.

Phylogenomic analysis of *L. platensis* allowed us to distinguish two major phylogenetic clades (I and II), but failed to clearly segregate strains following their habitats in terms of salinity or biogeography^20^. Such discrepancies have already been observed in other taxa, for instance for the widespread extremophilic bacteria *Salinibacter ruber*^22^. However, in this case, non-targeted metabolites analysis succeeded to segregate the strains, highlighting biogeographical metabolic signatures related to local adaptations. Recently, the same approach has suggested the existence of distinct chemical signatures between different ecotypes of the oceanic picocyanobacterium *Prochlorococcus marinus*, each being adapted to a specific niche characterized in terms of light intensity and temperature^23^.

Concerning the species *L. platensis*, the flexible part of the genome remarkably represents up to 55% of the genome of each strain, and differs from one strain to another, carries a large variety of genes (certain of them presenting regulatory and defensive functions)^20^, many of which present so-far unknown functions and could potentially be related to the specific capabilities of the strains in terms of physiology and ecology. One can presume that the genes found within this variable portion of the genome could be involved in *L. platensis* adaptation to local environmental conditions. These genes could indeed be responsible of carrying traits that allow the species to adjust to specific local conditions and support the maintenance of constant blooms. Ultimately, this local adaptation might give *L. platensis* an advantage to overpass other phytoplankton competitors within these environments.

In the present paper, we attempt to determine whether *Limnospira platensis* displays metabolic signatures specific to its different habitats, particularly brackish or alkaline-saline ecosystems, and question the impact of accessory gene repertoires on respective chemical adaptations. In this aim, we have investigated a wide panel of strains from different environments and maintained them in collections under identical culture conditions. We used two types of extraction and chemical analysis approaches, pursuing a large diversity of metabolites potentially involved in various adaptative traits. First, we applied a targeted approach, focusing on photosynthetic pigments and fatty acids known to be involved in adaptation to different environmental conditions (*e.g.*, light, salinity, and pH). Then, we performed an untargeted approach in parallel to obtain an overall chemical fingerprint of each strain and look for other potential differences.

## Material and Methods

### Strains and culture conditions

93 strains of *L. platensis* were studied. 89 strains were obtained from the Paris Museum Collection (PMC)^24^: 72 from two Camargue area, a humid region located in the Rhône delta (South of France) mostly consisting of marshland and brine lagoons, 13 from the lake Dziani Dzaha, an alkaline and thalassohaline lake located on the island Petite-Terre of Mayotte, three from lake Natron, an alkaline-saline lake located between Tanzania and Kenya, and one strain from historical collection with no environmental data available. Three additional strains were provided by other collections: *L. platensis* PCC 8005 and *L. platensis* PCC 7345 (Pasteur Culture Collection of Cyanobacteria, France) from unknown locations, and *L. platensis* SAG 85.79 (Culture Collection of Algae at Göttingen University, Germany) from the lake Natron. one strain provided by Spirulina Solutions (https://spirulinasolutions.fr/): Paracas R14 from a freshwater lake in Peru. All information about strains, sampling site properties, year of isolation, and genomic information is given in Supplementary Table S1. These 93 strains were maintained as monospecific but non-axenic cultures, growing in 30 mL of sensibly improved Spirulina medium^24^ at 24°C with a photon flux density of 30 µmol photon.m^-2^.s^-1^ under a 16:8-h light:dark photoperiod. When investigating the metabolite production of the various strains, careful attention was paid to maintaining cultures in the growing phase to keep the level of heterotrophic bacteria as low as possible. We assume it remains negligible comparing to cyanobacteria development, especially in terms of biovolume, biomass, and metabolite contents, and might not critically influence the present observation on *Limnospira* strain distinctions. Each month, 1/10 of each culture were used to inoculated new cultures. The remaining 9/10 were centrifuged (10 min, 3,220 g, 3 times). The supernatants were discarded, and the pellets were stored at -80°C for at least 3h, and then lyophilization was performed (-47°C, 0.03 mBar for 16h) (Labconco, Kansas City, MO, USA). Six successive cycles of culture-biomass sampling have been made and pooled. The six freeze-dried biomass samples of each strain have been mixed together and homogenized prior to the different chemical analyses for both pigments, fatty acids and untargeted metabolites.

### Pigments

Hydrophilic pigments of the 93 strains were analyzed by spectrophotometry according to Yéprémian et al.^25–26^. Ten mg of freeze-dried biomass were incubated with 10 mL of a saline buffer solution (NaCl 0.15 mol.L^−1^, KCl 27 mmol.L^−1^, Na_2_HPO_4_ 80 mmol.L^−1^, KH_2_PO_4_ 20 mmol.L^−1^, and Na_2_-EDTA 10 mmol.L^−1^; pH = 7.5) for 16h, at 4°C in the dark. Samples were then centrifuged (10 min, 3,220 g), and 1 mL of the supernatant was analyzed with a spectrophotometer (CARY-60, Agilent) at 565, 620, and 650 nm, corrected by subtracting the value recorded at 750 nm (< 0.01 absorbance unit). The values reported as concentration (mg.g^-1^) of C-phycocyanin and allophycocyanin (absence of phycoerythrin) were obtained with calculus according to de Marsac and Houmard^27^.

Lipophilic pigments of the 93 strains were analyzed by high performance liquid chromatography (HPLC). The complete procedure is detailed in supplementary material and methods. The relative abundance of each pigment (%) was calculated from its respective concentration in the sample (mg.g^-1^ dw for chlorophyll *a* derivative, and μg.g^−1^ dw for other pigments).

### Fatty acids

Ten mg of freeze-dried biomass were used for the extraction of fatty acids from 91 strains (all 93 except *L. platensis* PMC 1041.18 and PMC 1044.18). Prior to extraction, an internal standard (tricosanoic acid: 23:0) was added in each sample. Lipids were extracted according to the protocol of Bligh and Dyer^29^ as modified by Meziane et al.^30^. The complete procedure is detailed in supplementary material and methods. Fatty acids were identified with a mass spectrometer (Agilent 5977B GC/MSD) according to the comparison of retention times of commercial fatty acid standards (Supelco 37). We then reported the values as % of total FA and concentration (mg.g^-1^ dw).

### Mass spectrometry

The intracellular metabolite contents of 88 strains (all 93 except *L. platensis* PMC 1287.21, PMC 1288.21, PMC 1298.21, PMC 1300.21, and PMC 1302.21) were analyzed as previously described^31^. The complete procedure is detailed in supplementary material and methods. Intracellular metabolite contents were analyzed using an electrospray ionization hybrid quadrupole time-of-flight (ESI-QqTOF) high-resolution mass spectrometer (Compact, Brucker, Bremen, Germany) in the range 50–1500 *m/z*. Metabolite annotations were made from MS2 data by generating a molecular network for the comparison of fragmentation profiles using the MetGem software (version 1.3.6) with GNPS algorithm (supplementary figure S1).

### Statistical analyses

The MetaboAnalyst 5.0 platform (www.metaboanalyst.ca) was used to perform data matrix normalization according to *Pareto* and mean-centered and divided by the standard deviation for untargeted metabolite, and for fatty acid and pigment matrices, respectively.

All three matrices (annotated ions, fatty acids, and pigments) were combined in a single matrix further called “total metabolite matrix”. The weight of each group of variables was adjusted to avoid the analysis being influenced by the largest group. The weights are identical for the variables of the same group.

Clade based on presence/absence of coding gene sequences was obtained with dendrogram (Jaccard’s dissimilarity index, Ward reconstruction, Silouhette analysis to find the optimal number of clusters) using software R (version 1.4.1103).

Hierarchical clustering on principal components (HCPC) using Euclidian distance and a principal component analysis (PCA) were performed to find different groups of strain using software R (version 1.4.1103). An analysis of similarities (ANOSIM) was performed on the total metabolite matrix to test differences between factors. Multiple correspondence analysis (MCA) was performed to identify correlations between tested factors using software R (version 1.4.1103). For each tested factor, multiple factor analyses (MFA) and between class analyses (BCA) were performed simultaneously (BC-MFA, code available at https://github.com/Hubas-prog/BC-MFA) on the combined matrix, as describe in Michelet et al.^32^. The percentage of total inertia explained (TIE) by the instrumental variable was systematically calculated. A threshold based on the square cosine of the coordinates of the variables was applied for highlighting the most discriminating variables with a square cosine superior to 0.4 (cos^2^ > 0.4). A Monte-Carlo test was systematically performed to test the significance of the BCA ordination, using software R (version 1.4.1103). For each factor, principal component analyses (PCA) and heatmap analyses were performed on each of the three matrices using the MetaboAnalyst 5.0 platform.

Differences between each class of factor have been searched for each of the 215 molecules with Kruskal-Wallis tests followed by pairwise Wilcoxon tests. For total fatty acid means comparison, normality was present (Shapiro-Wilk test, p > 0.05) and homogeneity of the variances was not present (Bartlett test, p < 0.05), thus a Welch test was performed. All tests were run using software R (version 1.4.1103).

## Results

### Pigment composition

Fourty photosynthetic pigments were detected and quantified by HPLC-DAD in the 93 analyzed *L. platensis* strains: two phycobiliproteins (C-phycocyanin and allophycocyanin), 12 chlorophylls or derivatives (chlorophylls *a*, *d*, and pheophytins), and 26 carotenoids (five carotenes, four carotenoid glycosides, six xanthophylls, one unknown carotene, and ten unknown carotenoids) (Figure 1A; Supplementary Table S2). A mean of 27.9 ± 1.8 pigments was detected per strain, ranging from 20 to 32 for strain *L. platensis* PMC 1223.20, PMC 1278.20 and PMC 1279.20, and *L. platensis* PMC 1041.18, respectively (Supplementary Table 2). 12 pigments were detected in all strains: C-phycocyanin, allophycocyanin, chlorophyll *a,* chlorophyll *a* allomer, chlorophyll *a* epimer, two pheophytins *a*, β,φ-carotene, β,ε-carotene-like, zeaxanthin, echinenone, echinenone/cantaxanthin-like, and two unknown carotenoids (UC10, and UC9) (Supplementary Table S3). The most abundant pigments were C-phycocyanin (64.5 ± 3.1% of total pigment composition) and allophycocyanin (25 ± 1.6% of total pigment composition). The chlorophyll *a* allomer was the most abundant form of chlorophyll *a* in 90 out of 93 strains, representing 6.6 ± 2.3% of the total pigment composition. The three other strains (*L. platensis* PMC 1223.20, PMC 1278.20, and PMC 1279.20) had the usual form of chlorophyll *a* as the main chlorophyll *a*, representing 13.6 ± 2.3% of the total pigment composition (Supplementary Table 2). Carotenoids represented 1.19 ± 0.6% of the total pigment composition. The most abundant carotenoids were zeaxanthin (34.2 ± 5.5% of the total carotenoid composition), myxoxanthophyll (14.9 ± 5.6% of total carotenoid composition), β,β-carotene (9.9 ± 1.3% of total carotenoid composition), and UC9 (425, 448, 470nm) (8.7 ± 3.6% of total carotenoid composition) (Supplementary Table S2).

**Figure 1.**
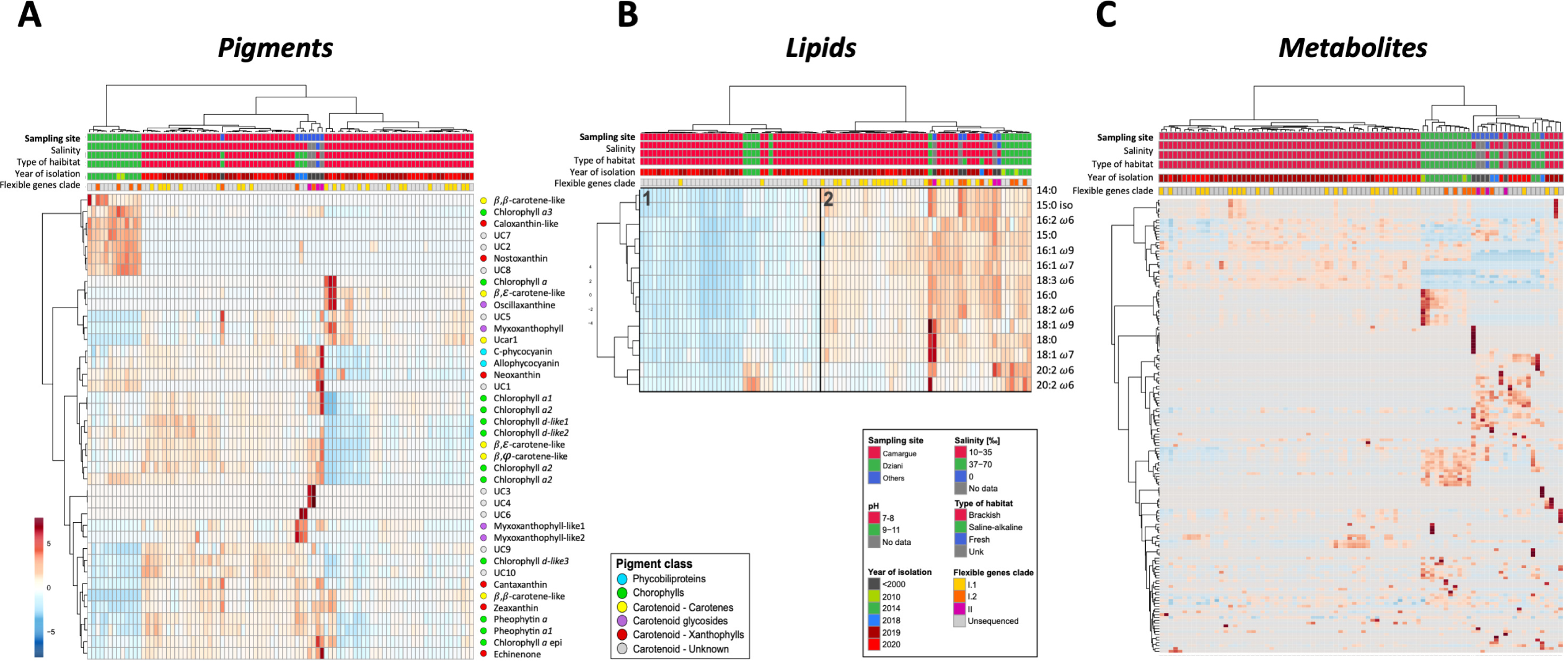
Heatmap analyses of pigment, lipid, and metabolite matrices. Heatmap analysis of the 40 pigment concentrations of the 93 analyzed *L. platensis* strains (normalization according to each feature) (A). Heatmap analysis of the normalized 14 fatty acid concentrations of the 91 analyzed *L. platensis* strains (B). Total fatty acid concentration (mg.g^-1^ dw) discriminates between 2 groups of lower (1) and higher (2) lipid contents (Welch test, p < 0.001). (C) Heatmap analysis of the 161 HPLC-MS/MS annotated ions of the 88 analyzed *L. platensis* strains, expressed in normalized relative abundances.

### Fatty acid composition

Fourteen fatty acids were detected and quantified by GC-FID and GC-MS in the 91 analyzed *L. platensis* strains. A mean of 52.32 ± 23.77 mg fatty acids.g^-1^ dry weight (dw) was quantified, ranging from 13.78 to 110.59 mg.g^-1^ for the strains *L. platensis* PMC 1259.20 and *L. platensis* PMC 894.15, respectively (Supplementary Table S3). Among the 14 fatty acids detected, three were saturated fatty acids (14:0, 16:0, and 18:0), two were branched fatty acids (15:0iso, and 15:0anteiso), four were mono-unsaturated fatty acids (16:1ω7, 16:1ω9, 18:1ω7, and 18:1ω9), and five were poly-unsaturated fatty acids (16:2ω6, 18:2ω6, 18:3ω6, 20:2ω6, and 20:3ω6). Five fatty acids represented > 90% of the total fatty acids: 16:0 (38.6 ± 1.8%), 18:3ω6 (24.6 ± 2.1%), 18:2ω6 (17.1 ± 1.5%), 16:1ω9 (6.2 ± 1.1%), and 16:1ω7 (5.6 ± 0.8%) (Supplementary Table S3). Interestingly, the relative quantities of these molecules within the different strains allow to most distinguish a group of “lower” and “higher” fatty acid-containing strains according to heatmap with hierarchical clustering with no obvious relation with the respective metadata of the strains (Figure 1B).

### Metabolite composition by LC-MS

A total of 2,851 ions were detected by HPLC-MS in the 88 analyzed *L. platensis* strains. A mean of 1,311.7 ± 175.1 ions was detected, ranging from 1,023 to 1,686 for strain *L. platensis* PMC 1044.18 and *L. platensis* SAG 85.79, respectively (Supplementary Table S4). 1013 ions (35.5%) exhibit an intensity superior to a threshold of 5,000 counts in single MS and were analyzed by HPLC-MS/MS. Among them, 221 ions (15.9% of fragmented ion, and 5.6% of total ions) have been clustered and annotated with MetGem and GNPS (*e.g.* phospholipids, saccharides, peptides, …) and 25 have been annotated at the molecular levels (*e.g.* ergothioneine, glutathione, biopterin, nicotinamide adenine dinucleotide) (Figure 1C; Supplementary Table S4), whereas other were annotated according to MS/MS fragmentation pattern similarity revealed by molecular networking (Supplementary Figure S1). Analyses of the 2,851 total ions and the 221 annotated ions showed similar results (Data not shown). We further focused only of the 221 annotated ions (Figure 1C).

### Groups of strains

Based on all dataset from both fatty acids, pigments and annotated ions, a complete matrix further called “total metabolite matrix” was assembled for performing global metabolite analyses (see Materials and methods). Considering this total metabolite matrix, a multiple factor analysis (MFA) combined to a hierarchical clustering on principal components (HCPC) was performed considering the effect of the sampling site factor and highlighted three distinct groups of strains (Figure 2A and 2B, respectively). Remarkably, those three groups that were statistically different (Figure 2A, ANOSIM p < 0.001) and also similarly discriminate regarding both fatty acids, pigments and annotated ions, when considered separately (supplementary Figure S2A-C), were overall matching with the sampling location factor (Figure 2C). Indeed, the strains isolated from different sampling sites were clearly clustered into different groups: Camargue region in group 1, Lake Dziani Dzaha in group 3, and another sampling site in group 2 (Figure 2B). Indeed, these analyses showed that strains coming from the lake Natron, Peru, and from the three unknown locations were overall grouped together in a single group 3 (Figure 2A-B). Thus, those strains of group 3 were further considered together in a site group named “Others”. According to the multiple correspondence analysis (Figure 2D), the correlation between these three groups (1=“Camargue”, 3=“Dziani” and 2=“Others”) and the different environmental factors was investigated. Group 1 mostly correlated to the range of pH of 7-8, salinity of 10-35‰, brackish habitat, and years of isolation 2019 and 2020, while Group 3 correlated to the range of pH of 9-11, salinity of 35-70‰, alkaline-saline habitat, and years of isolation 2010 and 2014, and Group 2 with other pH, salinity, habitat, and year of isolation prior to 2000 (Figure 2 C-D). The three “sampling site” groups highlighted by the HCPC were used for further analysis regarding their respective metabolite signatures.

**Figure 2.**
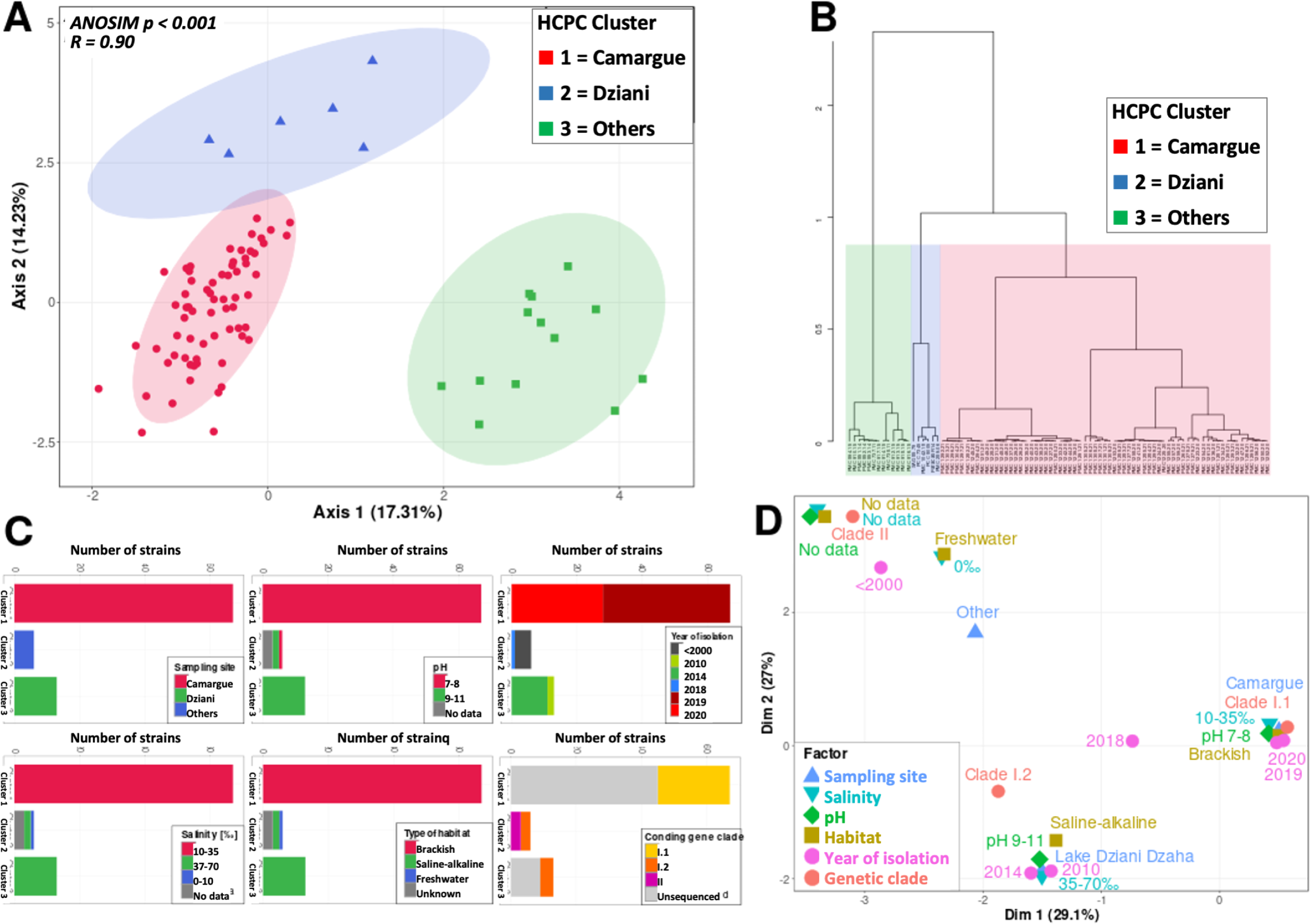
Principal component analysis (PCA) (A) represents the total matrix metabolite composition of the strains regarding the groups given by the hierarchical clustering on principal components (HCPC) (B). Characterization of each group regarding the sic tested factors (C) and the multiple correspondence analysis (D).

### Sampling sites

MFA and BCA analyses were performed considering the three groups that were previously separated by HCPC and clearly correspond to the different sampling sites (Figure 3A-B). The primary axis of the BCA analysis accounted for 63.1% of the variability. It primarily distinguished the *L. platensis* samples taken at Dziani (group 3) from the others, based on their chemical composition (Figure 3B). The second axis explained 37% of total inertia and mainly opposed HCPC group 2 from all other strains (Figure 3B). Among fatty acids, pigments, and annotated ions, 31 most discriminant variants were observed and present a cos^2^ > 0.4 (Figure 3C): two fatty acids, nine pigments, and 20 annotated ions (26 related to axis 1, and five to axis 2). In total, 126 molecules showed significative difference between at least two groups (Kruskal-Wallis test, p < 0.001). The two fatty acids were the PUFAs with the longest carbon chains, 20:2ω6 and 20:3ω6 that were significantly more present in the strains from lake Dziani Dzaha in concentration and proportion of total fatty acids (Figure 4, Kruskal-Wallis test, p < 0.001). Based on fatty acids only, two groups of 42 and 49 strains have been discriminated by hierarchal clustering, not related to any of the environmental factor (Supplementary Figure 2B). They significantly differed by the total fatty acid biomass (30 ± 8 mg.g^-^and 71 ± 15 mg.g^-1^, respectively).

**Figure 3.**
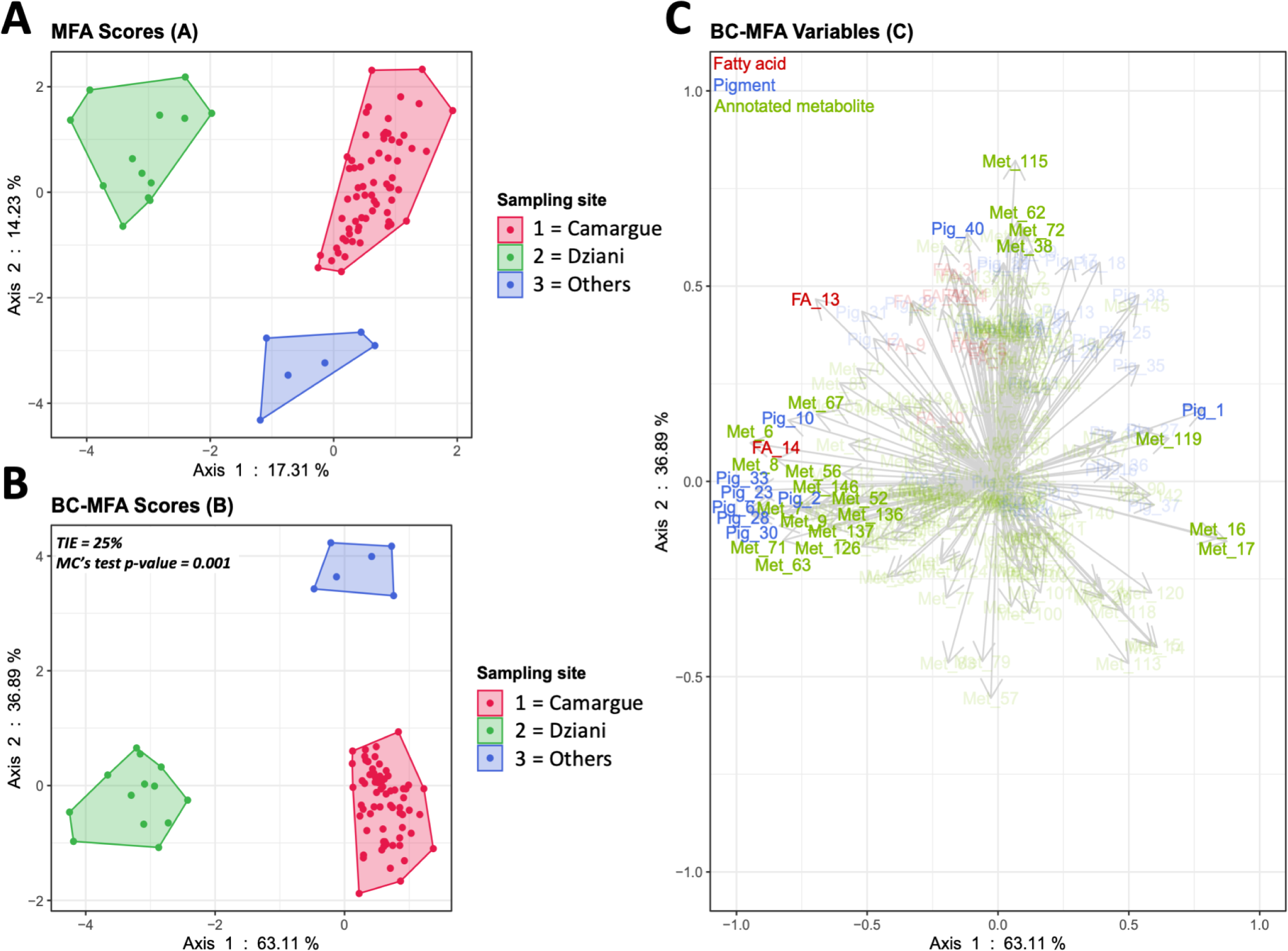
Multiple factor analysis (MFA: A), and between class analysis with a cos^2^ threshold of 0.4 (BCA: B, cos^2^ plot: C) based on the chemical composition of the strains. The tested factor is the sampling site.

**Figure 4.**
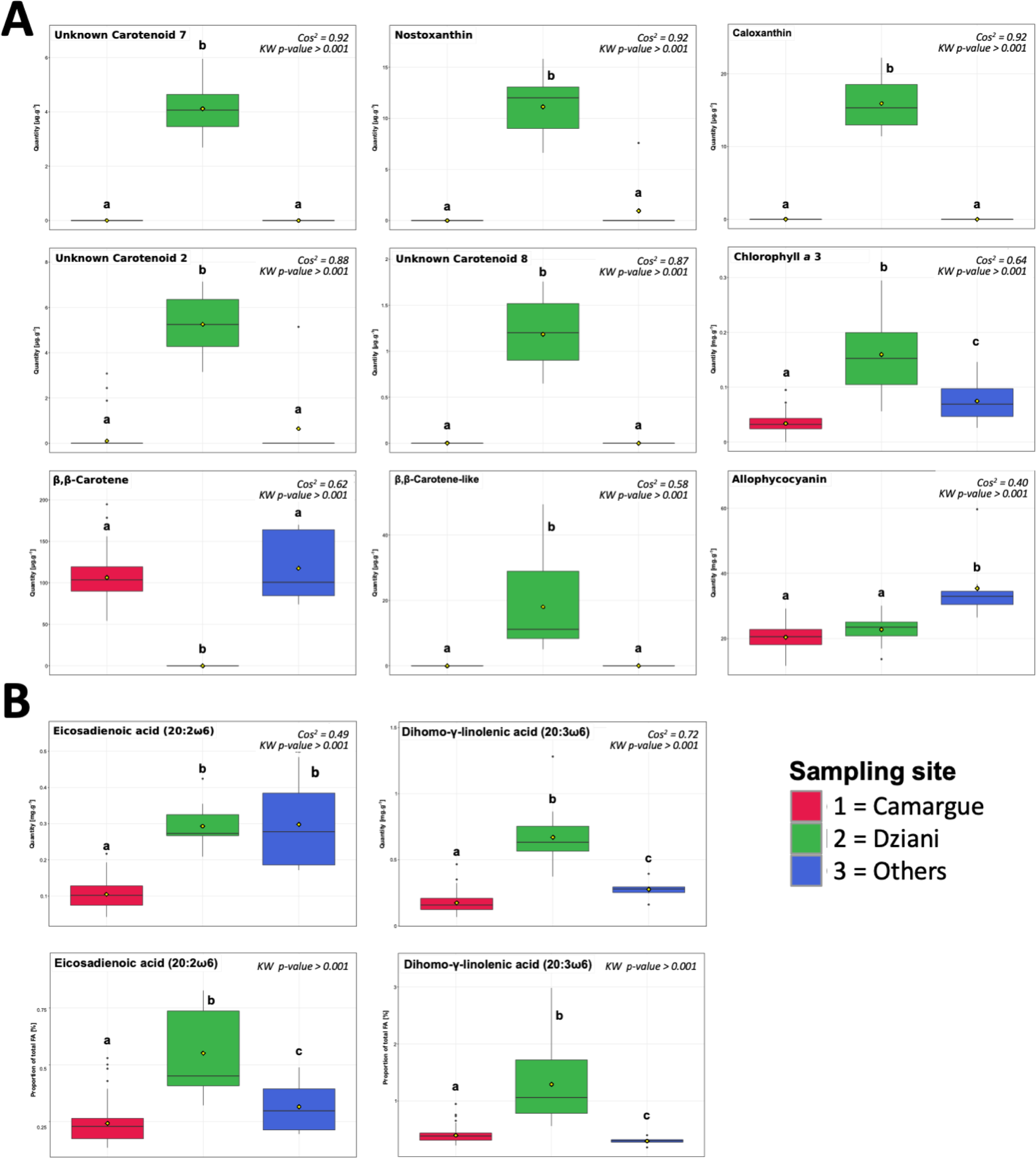
Most discriminant fatty acids and pigments (cos^2^ > 0.4) were compared between sampling sites (Kruskal-Wallis test, pairwise Wilcoxon tests). Pigment concentrations are given in mg.g^-1^ dw for phycobiliproteins and chlorophylls a, and µg.g^-1^ dw for other pigments (A). Fatty acid concentrations are given in mg.g^-1^ dw, and proportions in % of total fatty acids (B).

Six of the pigments were only present in the strains from lake Dziani Dzaha (Figures 1A and 4, Kruskal-Wallis test, p < 0.001). Among them, three were recognized as resembling caloxanthin and nostoxanthin and an analog of β,β-carotene, and three were unknown carotenoids. The strains from lake Dziani Dzaha did not possessed the usual form of β,β-carotene, that was only present in the strains from Camargue and other locations. The allophycocyanin were significantly more present in the strains from other locations (Figure 4, Kruskal-Wallis test, p < 0.001). Among the 20 most discriminant annotated metabolites (Figure 5), 13 were significantly more abundant in the strains from lake from lake Dziani Dzaha, including four saccharides (MW = 254.1022, 596.2184, 684.2349 Da, and sucrose), three little peptides of 2 or 3 amino acids (MW = 373.2214, 359.2057, and 260.1376 Da), two phospholipids (MW = 467.304, and 479.3041 Da), one lipid (MW = 455.3042 Da), and two unsaturated fatty acid glycerol annotated as monoolein and alpha-linolenoyl-glycerol (Kruskal-Wallis test, p < 0.001). Three molecules were significantly less abundant in the strains from lake Dziani Dzaha: two saccharides (MW = 285.1065, and 268.0803 Da), and S-methylglutathione (Kruskal-Wallis test, p < 0.001). Four molecules were significantly more present in the strains from other locations: two peptides (MW = 359.1869, and 373.1986 Da), one fatty acid ester (MW = 588.3765 Da), and oxidized glutathione (Kruskal-Wallis test, p < 0.001).

**Figure 5.**
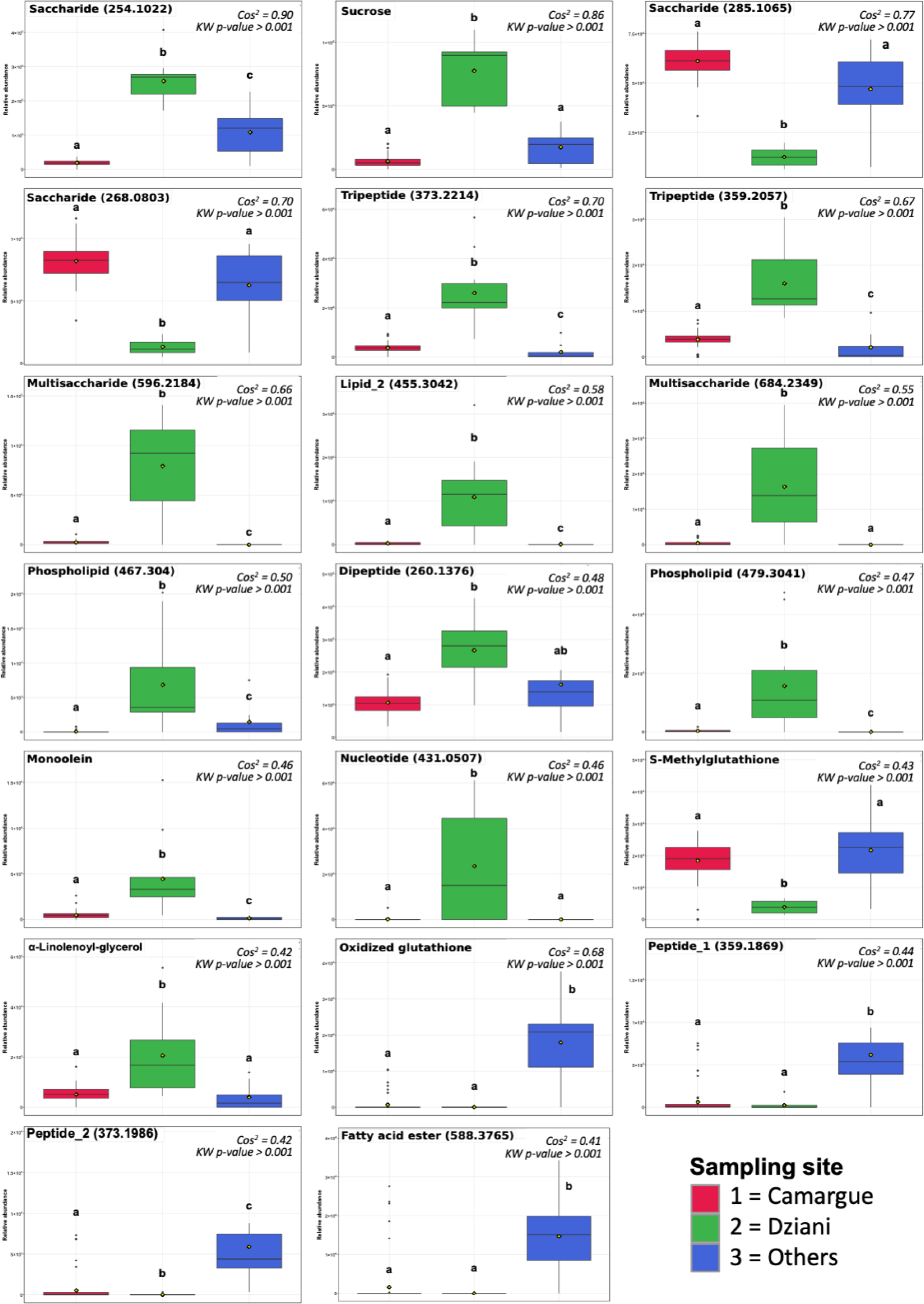
Relative abundances of most discriminant annotated ions (cos^2^ > 0.4) were compared between sampling sites (Kruskal-Wallis and pairwise Wilcoxon tests).

### Gene content

Among the 93 *Limnospira* strains presently investigated, 33 of them present already available genomic information (supplementary table S1) that could explored for the detecting distinctive genetic traits. that could be related to the clustered strains according to their chemical signatures. Interestingly, the analysis of the presence/absence of coding genes perfectly match with distance pattern between the strains determined according to the core-genome sequence comparison, by distinguishing two main groups of strains (I and II) (supplementary figure S3). The first group is divided into two subgroups (I.1 and I.2); the clade I.1 contains 23 strains from the Camargue region, and the clade I.2 four strains from Lake Dziani Dzaha, two strains from Lake Natron (*L. platensis* PMC 1042.18 and SAG 85.79, and the strain *L. platensis* PCC 8005. The clade II contains the last three strains (*L. platensis* PCC 7345, PMC 289.06, and Paracas R14).

Interestingly, these three coding gene clades (I.1, I.2 and II) can also be clearly discriminated according to both MFA and BC-MFA analyses (Supplementary Figure S4A-B). The chemical composition of all three clades was mostly separated by the first axis of the MFA and the BC-MFA, while all clade II were separated from clades I.1 and I.2 by the second axis of the BC-MFA. Among fatty acids, pigments, and annotated ions, 18 most discriminant molecules were observed on BC-MFA variable plot to present a cos^2^ > 0.4 (Supplementary Figure 4C): seven fatty acids, seven pigments, and four annotated ions (17 related to axis 1, and one to axis 2). In total, 63 molecules showed significant difference between at least two groups (Kruskal-Wallis test, p < 0.01). The seven fatty acids, as well as the seven pigments, were less abundant in clade I.1 (Supplementary Figure S5, Kruskal-Wallis test, p < 0.01). Among the four most discriminant annotated ions, two saccharides (MW = 536.1602 and 430.133 Da) were significantly more abundant in the strains of clade I.1. The two other annotated ions were significantly more abundant in the strains of clade II: oxidized glutathione and a fatty acid ester (MW = 588.3765 Da) (Supplementary Figure S5, Kruskal-Wallis test, p < 0.01).

## Discussion

In the present work, we have developed an original approach based on the investigation specific adaptation of the different chemotypes to their original environmental conditions, through the observation of their regular chemical traits. Thus, we hypothesized that the original chemical traits of these different genotypes might offer them remarkable adaptative features that have been shaped by the specific conditions of their respective originating environments.

Indeed, analyses of the pigment, lipid, and small metabolite contents of various *Limnospira platensis* strains cultivated under identical laboratory conditions provide unprecedented data that offers a plain answer to the foremost question: is the diversity of these molecules driven by selective pressures on *Limnospira platensis* (clade I or II)^20^ and/or by the environment from which the strains are isolated? Our results clearly show that the sampling location that offers diverse environmental conditions (*e.g.* salinity, pH, light, or temperature) constitutes the variables explaining the distribution of *Limnospira platensis* in three distinct chemotype groups. The molecules that most discriminate these three groups of the *Limnospira platensis* strains studied, under the standard conditions of lab cultures, are saccharides, fatty acids, peptides, photosynthetic pigments, and antioxidant molecules. Interestingly, these molecule enrichments might constitute physiological adaptation traits to the local conditions encountered in their respective sampling environments, potentially in relation to local salinity, light and oxidative stress. These points are then critically debated in the following discussion.

### Osmoregulation and salinity resistance

The ponds of the Camargue in France and Lake Dziani Dzaha in Mayotte are environments with different salinities. In the brackish ponds of Camargue, close to the Mediterranean Sea, salinity varies between 12-35‰. Lake Dziani Dzaha is a thalassohaline lake. Here, salinity is much higher, reaching up to 70‰^33^. Strains from Lake Dziani Dzaha have greater sugar diversity, have more long-chain polyunsaturated fatty acids compared with strains isolated from other habitats under our experimental conditions. They are also the only ones to produce nostoxanthin and caloxanthin. This specific chemical signature of strains from Lake Dziani Dzaha testifies to the adaptation of the *Limnospira platensis* species to resist to higher salinity in this specific environment.

In cyanobacteria in general, several families of molecules are involved in osmoregulation and detoxification of stress due to high salinity^34–35^. Cyanobacteria produce various sugars (*e.g.* glucose, sucrose, trehalose or glycine betaine) to prevent the osmotic stress caused by high salinity^36^. In *Synechococcus*, long-chain polyunsaturated fatty acids help combat the oxidative stress caused by high salinity as they appear to be produced to a greater extent with increasing salinity^37^. Interestingly, it has also been shown that certain carotenoids, nostoxanthin and caloxanthin, are produced by a bacterium of the genus *Sphingomonas* that appears resistant to high salinity^38^. In addition, the nostoxanthin and caloxanthin-producing bacterium *Sphingomonas nostxanthinifaciens* may have a role in reducing the amount of reactive oxygen species (ROS) in *Arabidopsis thaliana* when the two organisms are co-cultured in high salt concentrations^39^. Taken together, these finding support the idea that sugar and carotenoid molecules might support high salinity adaptation process in *Limnospira platensis*.

### Light intensity

Facing high light, cyanobacteria modify chlorophyll, phycobiliprotein, and carotenoid composition to improve electron transfer and manage excess energy^40–42^. In *Synechococcus*, various carotenoids are crucial for high light survival. Strains lacking carotenoid biosynthesis genes (*cru* genes) have lower growth rates and higher ROS levels compared to carotenoid-producing strains^40^.

The different *Limnospira platensis* strains relied on a wide diversity of chlorophylls (n = 5-12) and carotenoids (n = 13-18) that could accordingly be involved in photo-pigment adaptation. One of the most striking results is the presence of six exclusive carotenoids that appear exclusively in strains from Lake Dziani Dzaha. This lake has a mean surface irradiance of 2,000 µmol photon.s^-1^.m^-2^, which decreases by 99% in the first 20 cm of surface water to around 20 µmol photon.s^-1^.m^-2^.^33^ *Limnospira* living in the photic zone of the lake are therefore subjected to both high and strong variations in light intensity over shallow depths. We found 18 carotenoid pigments that have an unknown chemical structure (putative analogues of other pigments, n = 7, or totally unknown pigments, n = 11). This observation illustrates the ability of *Limnospira* to diversify its pigment content in response to a given environment. The great diversity of pigments produced by *Limnospira platensis* was shown by the characterization of up to 48 hydrophobic pigments (22 carotenoids and 26 chlorophylls) by LC-APCI-FT-ICR-MS^43^, some of which could very likely correspond to those unknown pigments observed in our study.

### Oxidative stress

In a general manner, both high salinity, light and UV exposures are interdependently involved in ROS production and subsequent cellular oxidative stress^44^. In addition to primary mechanisms listed above, aiming at reducing the adverse effect of oxidative stress, cyanobacteria also produce a large variety of other molecules (other than carotenoids and long-chain polyunsaturated fatty acids). These molecules are specifically involved in buffering ROS production and combating oxidative stress deleterious effects, such as ergothioneine, glutathione, biopterins and their derivatives^14,45–46^. The cellular extracts of all *Limnospira platensis* strains presently investigated contained significant amounts of each of these molecules, that might constitute a set of molecular resource for this organism to deal with remarkable local cellular stressors.

### *Limnospira platensis* presents a peculiar ecological strategy

Cyanobacteria have colonized many types of habitats (*e.g.* oceanic waters, fresh waters, alkaline waters, benthic or pelagic, terrestrial, symbiotic)^2^. Aquatic species have for example developed different ecological strategies to adapt to their environment. Some cyanobacteria live in diversified communities of various species being concomitant or consecutive along the proliferative events. These include benthic cyanobacteria, which form biofilms (*e.g. Phormidium*, *Lyngbya*, *Oscillatoria*)^47^, and pelagic cyanobacteria, which can form blooms (*e.g. Microcystis*, *Aphanizomenon, Raphidiopsis, …*)^48–50^. The genomes of these cyanobacteria are remarkably large (> 5Mb) and contain many BGCs^51–53^, producing various bioactive secondary metabolites (including cyclic peptides, lipopeptides, alkaloids, macrolides, polyketides, etc.) that enable cyanobacteria to interact with their microbial community (*e.g.* anti-grazing, anti-algal, communication blocking, competition for resources)^5,54^. However, the production of these specific molecules has a high energy cost for the producing cells, as they are involving large gene sets and the synthesis of many enzymes^55^. In contrary, some marine picocyanobacteria (*e.g. Prochlorococcus*, *Synechococcus*) are exclusively dominating one type of habitat (*i.e.* the worldwide oligothrophic seas) and exhibit a unique diversity of photosynthetic pigments, allowing them to exploit a wide range of light micro-niches^56–57^. The genomes of these cyanobacteria are remarkably small (1.5-2.5Mb) and contain no BGCs^51,58^. In these cyanobacteria, adaptation to an ecological niche is encoded in both core and flexible parts of the genomes^59–60^.

For *Limnospira platensis*, we denote a potential third strategy. *Limnospira platensis* genomes are relatively large, ranging between 5.5 and 6.5Mb^20,61^, being comparable to the genome size of community-dwelling cyanobacteria proliferating in eutrophic waters. Yet, they remarkably contain no or only one BGC of secondary metabolites (potentially involved in the biosynthesis of cyanobactins called Arthrospiramides), as do the specialized marine picocyanobacteria *Synechococcus* and *Prochlorococcus*^51,62–63^. Unlike marine *Synechococcus* and *Prochlorococcus*, *Limnospira platensis* proliferates in many different habitat types: freshwater, brackish water, alkaline water, and saline alkaline water^20,64^, and *Limnospira platensis* strains belonging to the same phylogenetic clade do not share similar habitats^20^. Moreover, in lakes where phytoplanktonic communities are dominated by *Limnospira platensis*, this species frequently generates persistent proliferations. These blooms are often accompanied by minimal surrounding diversity (except de picoeukaryote *Picocystis salinarum*)^21,63^, serving as a distinctive indicator of extreme environments (*e.g.*, hypersaline conditions, high light coupled with high turbidity, elevated temperatures, high pH levels, and high sulfur contents). In such environments, only a limited number of microorganism clades can develop. This absence of surrounding microbial diversity, comprising potential competitors, could explain the absence of BGCs in the *Limnospira platensis* genomes, as it has fewer phototrophic organisms with which to interact through allelopathic compound production, as various cyanobacteria from high diversity freshwater temperate lakes do^51^. Interestingly, such an ecological strategy could support the high biomass production observed both *in situ* and *in vitro*, as lab culture conditions present many similarities (in terms of both biotic and abiotic factors) with those of the extreme environment in which *Limnospira* proliferates. Indeed, it has been shown by transcriptomics that the essential transcribed functions expressed *in vitro* are related to primary metabolism (metabolism of sugar, fatty acids, nitrogen, CO_2_, photosynthesis, and antioxidant processes)^64^.

Interestingly, the genomic investigation of *Limnospira platensis* strains shows that both well-conserved (core) and flexible (pan) genome parts discriminate strains according to their respective origin locations (supplementary Figure S3). Thus, one can suppose that either common point or InDel mutations on key metabolic genes of the core-genome or variation in the flexible gene content of *Limnospira platensis* genomes may support peculiar adaptations to local environmental conditions. Interestingly, this latter flexible part of the genome is remarkably rich in transposable and regulatory elements and contains various genes, most of which have as yet unknown molecular functions, and may dynamically support evolutionary exaptation processes such as ecological niche extension due to higher stress resistance aquisition^20^. So far, the relationship between small metabolite content (*i.e.,* redox agents, saccharides and pigments/carotenoids) - chemotype - and niche partitioning (*i.e.,* ROS scavenging, osmoregulatory, and photoprotective molecular traits) - ecotype - still remains largely undetermined due to the lack of distinctive genetic markers associated with the various anti-oxidant, osmolyte and pigment production. Such assumptions now require further experimental validation.

## Conclusion

Taken together, these ecological and evolutionary concerns offer new comprehension of the mechanisms that support the remarkable success encountered by *Limnospira* in natural as well as lab or industrial environments. One step further would be to try to experimentally identify the genetic and phenotypic traits that are defining the different *Limnospira* ecotypes and to determine whether *Limnospira* clade I.1, I.2 and II genotypes can colonize similar ecological niches, as this taxon exhibits ubiquitous geographic distribution, or represents already well-distinct ecologically and functionally distinguishable taxa.

Also, key questions remain on the understanding of the mechanisms that allow *Limnospira in situ* and *in vitro* blooms to avoid density-dependent regulations of its proliferation by cyanophages, as those encountered by other blooming cyanobacteria^65^. Such adaptative mechanisms would help explain how *Limnospira platensis* can form permanent blooms in many environments. To this end, two distinct hypotheses can be further tested: either the *Limnospira platensis* genome presents a mechanism to protect itself from phage lyses, or the fraction of the population potentially lysed by phages would be replaced by rapidly dividing *Limnospira platensis*.

## Author Contributions

Conceptualization, T.R., J.-P.C., L.V., C.B. and B.M.; methodology, T.R., C.H., M.C., C.Y., B.M.; investigation, T.R., C.H., T.M.; resources, C.D.; writing-original draft preparation, T.R., C.B. and B.M.; writing—review and editing, all authors. All authors have read and agreed to the published version of the manuscript.

## Acknowledgments

We are grateful to the anonymous reviewer for their valuable comments and suggestions, which have helped improve the quality of our manuscript. We would like to thank the UMR 7245 MCAM, Muséum National d’Histoire Naturelle, Paris, France for laboratories facilities and Algama for funds. The MS spectra were acquired at the Plateau technique de spectrométrie de masse bio-organique, Muséum National d’Histoire Naturelle, Paris, France. The authors thank the Pasteur Collection of Cyanobacteria, the Culture Collection of Algae at Göttigen University, and Gilles Planchon for sending strains.

